# Distribution of carbon monoxide-oxidizing microorganisms along a chronosequence on Piton De La Fournaise volcano

**DOI:** 10.1101/2025.11.28.691264

**Authors:** Constance Wildbur, Robin A. Dawson, Shamik Roy, Claudine Ah-Peng, Mikk Espenberg, Marcela Hernández

**Affiliations:** School of Biological Sciences, University of East Anglia, Norwich, NR4 7TJ, UK; Chair for Forest Zoology, Technische Universität Dresden, Tharandt 01737, Germany; UMR PVBMT, Université de la Réunion, 97410, Saint-Pierre, La Réunion, France; OSU-Réunion, Université de la Réunion, 97400, Saint-Denis, La Réunion, France; Department of Geography, Institute of Ecology & Earth Sciences, University of Tartu, Tartu, Estonia

**Author notes:** Corresponding author: Marcela Hernández, School of Biological Sciences, University of East Anglia, Norwich, NR4 7TJ, UK.

**Keywords:** carbon monoxide dehydrogenase, *cox* genes, metagenome-assembled genome, soil microbes, volcano

## Abstract

Volcanic soils provide a unique environment for studying microbial colonization and succession due to their extreme conditions and distinct geochemical profiles. This study focused on carbon monoxide (CO)-oxidizing microbial communities in volcanic soils of varying ages at Piton De La Fournaise (Réunion island). Soil samples from three sites were analyzed to assess microbial community structure using 16S rRNA gene sequencing and metagenomic analysis to identify functional genes involved in CO oxidation. The activity of CO oxidizing microbes in soils was measured. Phylum-level analysis showed increasing Acidobacteriota and Chloroflexota, decreasing Actinomycetota and Bacteroidota, and stable Pseudomonadota, while class-level patterns included rising Alphaproteobacteria and Acidobacteriia, with Ktenobacteria emerging in the oldest soils. CO dehydrogenase-related genes were found in 17 metagenome-assembled genomes across all sites. CO-oxidizing microbes were present across soil ages, with detectable activity in the younger soils and greatest activity in the oldest, suggesting that these microbes actively use CO as an energy source even in soils with primary vegetation, contrary to general understanding. The findings highlight the intricate dynamics of microbial succession in volcanic soils and challenge conventional expectations about community complexity over time. Understanding pioneer communities elucidates soil restoration processes, which will become critical when countering anthropogenic soil degradation.

## Introduction

Volcanic sites provide a model for analyzing how soil is formed through microbial interactions starting off with important pioneer microbes that are capable of surviving in extreme environments (Wubs *et al*. 2016, Fantom et al., 2025). The young volcanic soils can be classified as extreme environments due to the very low amount of organic matter (King 2003a; Fujimura *et al*. 2011).

These pioneer microbes display physiological and functional traits, which significantly influence nutrient cycling, thereby allowing soil succession to occur (King 2003a). Understanding how these pioneer microbes can inhabit these sites and influence succession helps us to understand more about effective soil restoration, which is becoming more important due to the anthropogenic degradation of the environment. The lack of organic matter in young volcanic deposits leads microbes to look elsewhere for potential energy and carbon sources. It is known that volcanoes produce trace amounts of reduced gases such as carbon dioxide (Di Muro *et al*., 2016), carbon monoxide (CO), hydrogen sulfide, hydrogen, methane, and through metagenomic functional analysis on soil samples collected from various volcanoes worldwide, genes which allow microorganisms to use CO as carbon and energy sources have been identified (Shepherd 1925; Hernández *et al*., 2020a; Robb and Techtmann 2018). Therefore, the development rate of these ecosystems could depend on the microbial utilization of trace gases as energy sources (King 2003a).

Carbon monoxide dehydrogenase (CODH) is a key enzyme in catalyzing the oxidation of CO in microorganisms. The gene clusters encoding CODH comprise of various subunits with the Mo-Cu-containing active site found in the large subunit, encoded by *coxL*. Various genes involved in the CODH gene cluster have been identified in bacteria from the phylum Acidobacteriota, Actinomycetota, Chloroflexota, Pseudomonadota, and Bacillota (Hernández *et al.,* 2020a; Dunfield and King 2004; Schübel *et al.,* 1995; Dawson *et al.,* 2025), reflecting the ecological adaptation of CO-oxidizing microorganisms to different environments.

This study investigates the pioneer microbial communities and how they change over time by analyzing a chronosequence pathway from volcanic samples from Réunion Island (Indian Ocean). Réunion island is a five-million-year-old basaltic volcanic edifice composed of the inactive Piton des Neiges and active Piton de la Fournaise shield volcanoes located southeast of the island. Piton de la Fournaise is known to be one of the most active volcanoes in the world (Morandi *et al.,* 2016). The climate of Piton de la Fournaise is humid and tropical, with an annual mean temperature range from 18 to 25°C and the driest months of precipitation between 75 and 400 mm (Jumeaux *et al.,* 2011).

Therefore, the lower part of Piton de la Fournaise is covered by tropical rainforests (Strasberg 1994). The lava flows can reach various elevations and sometimes even the ocean. Any flows less than 1000 years old have been observed to have very thin topsoil (Meunier *et al.,* 2010). Newly exposed volcanic deposits harbour diverse microbial communities despite harsh conditions such as low pH and minimal organic matter, comparable to other extreme terrestrial environments (Gomez Alvarez *et al.,* 2016).

Notable patterns emerged when investigating microbial succession on volcanic deposits, highlighting the critical role of suitable substrates for energy and biosynthetic metabolism, with high microbial abundance and diversity influenced by environmental gradients such as increased organic carbon and vegetation presence (Weber and King 2010; King 2003a). At Llaima Volcano in Chile Chloroflexota, Verrucomicrobiata, and Planctomycetota were found to dominate the younger soils, while Pseudomonadota and Acidobacteriota prevailed in the older sites (Hernández *et al.,* 2020b). The microbial populations in particularly vegetated, young soils exhibited versatile behaviors, as they contained the genetic potential to oxidize CO, hydrogen, or methane as energy sources, indicating facultative chemolithoautotrophic or mixotrophic capabilities (Hernández *et al.,* 2020a).

Studies conducted on CO oxidizers within volcanic environments offer valuable insights into their widespread presence and role in contributing to the development of complex microbial communities during soil formation. For instance, research at Llaima Volcano in Chile has uncovered the presence of CO oxidizers across various soil ages by recovering metagenome-assembled genomes (MAGs) from this environment containing the *coxL* genes (Hernández, et al., 2020a). Moreover, CO oxidizer abundance and diversity tend to increase with vegetation and organic matter concentrations (King 2003a; Weber and King 2010), similar to other microbes found in these communities, although the impact of CO oxidation may be negligible in vegetated environments (King and Weber 2008). Despite extreme conditions, volcanic deposits support a surprisingly diverse microbial community (Guo *et al.,* 2014; Byloos *et al.,* 2018), with aerobic CO oxidation being documented across numerous bacterial lineages revealing novel functional capacities within these environments (Dunfield and King 2004; Dawson *et al.,* 2025). Given that Piton de la Fournaise is an island volcano with strong elevational gradients, we aimed to investigate whether CO-oxidizing groups such as Pseudomonadota, Acidobacteriota, and Ktedonobacteria follow similar succession patterns to those reported from continental volcanic systems. We also sought to determine whether CO oxidation remains an active metabolic process in older soils where vegetation and organic carbon inputs have become established. We hypothesized that microbial community complexity and abundance increase with soil age and that CO oxidizing microbes would be predominantly active in younger soils, where organic matter is limited.

## Materials and Methods

### Sample collection and physicochemical properties determination

Soils were sampled in November 2022 at three different locations of the volcano: Piton de Bert (PDB) (2242 masl, 21.2788831 S, 55.6980607 E), Mare Longue (ML) (282 masl, 21.3512651 S, 55.7392276 E) and Coulée de lave (CDL) (124 masl, 21.2866284 S, 55.7957900 E) (Fig. 1), which were formed by eruptions in 1401, 1559 and 2007, respectively (Albert et al., 2020). Soil samples from the youngest site (CDL) were collected from the rock crevices. Three distinct vegetation types are represented at these three sites. PDB hosts subalpine vegetation composed of endemic shrubs such as *Erica reunionensis* E.G.H. Oliv., which are of small stature (<2 m). ML represents a rich native primary lowland forest with a canopy height of about 20 m. CDL hosts pioneer lowland vegetation, sparsely vegetated and composed of the lichen *Stereocaulon vulcanii* (Bory) Ach., two dominant moss species (*Campylopus aureonitens* (Müll. Hal.) A. Jaeger and *Polytrichum commune* Hedw.), and the fern *Nephrolepis abrupta* (Bory) Mett. Two cm of topsoil was removed at each site, and three random samples were taken in the pattern of a triangle for each site. Samples were shipped in Ziploc bags and then stored at 5 °C in the laboratory. Air and soil temperatures were recorded *in situ* using a temperature logger (Comet Systems Ltd., Rožnov pod Radhoštem, Czech Republic). Measurements may vary due to local conditions including canopy cover, elevation, and cloud cover.

**Figure 1.**
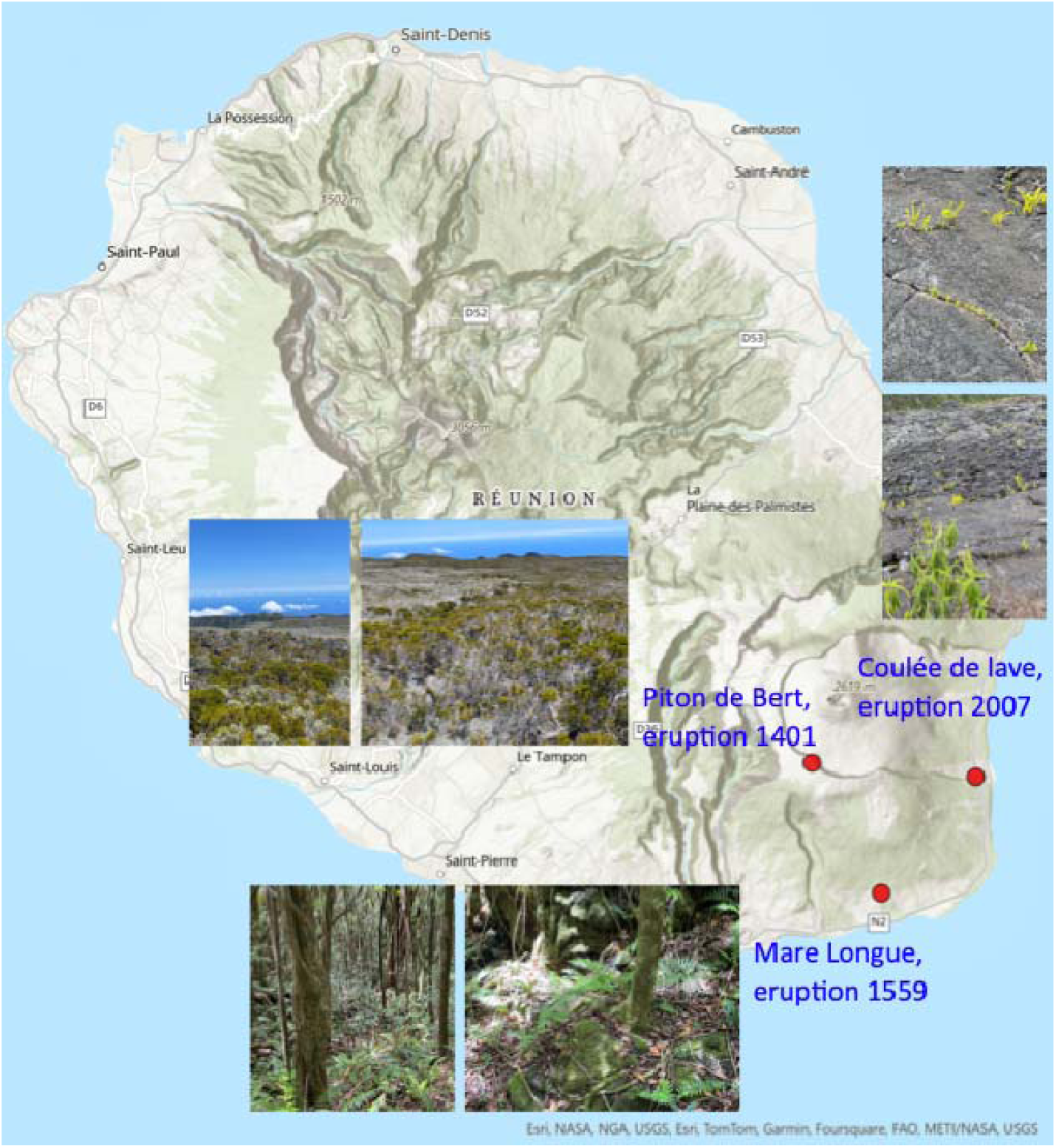
Map of the studied lava flows of Piton de la Fournaise volcano on Réunion Island.

pH was measured by mixing soil samples with distilled water in a 1:10 ratio (w/v), shaking samples at 180 rpm for 5 minutes in an orbital shaker and allowing to rest for 2 hours before measuring with a FiveEasy pH meter (Mettler Toledo). Soil moisture content was determined by recording the weight of the empty container and weight of soil before incubation, storing samples overnight in an oven at 100 °C, and then recording dried soil weights. Soil moisture content was then calculated as a percentage (Table S1).

### CO uptake by soil microbes

For all samples, 12 g of soil was sifted to remove rocks and plant matter and added to a 120 ml serum vial. CO was added from a 5000 ppm (0.5% in N_2_) stock solution to a final concentration of 100 ppm. CO concentrations were recorded using an Agilent Technologies 7890A Gas Chromatograph according to Dawson *et al.,* (2025). Briefly, sample gases were separated using an HP-Molesieve PLOT column (Agilent). CO was converted to methane by hydrogenation reaction on a nickel catalyst (350 °C) before detection by a flame ionization detector. Measurements were taken until CO consumption was no longer detected, and CO consumption rates were calculated from the linear decline in headspace CO over time, using the slope of a linear regression of CO concentration (µmol) versus incubation time (min), which was then converted to µmol h⁻¹ and normalised per gram of soil. Standards were prepared in N_2_-flushed 120 ml vials from 100-1 ppm CO. To ensure the activity in the observed vials was biological, control samples were made by autoclaving the soil for 60 minutes and leaving residual CO to diffuse out in a fume hood. The activity was measured by the same method indicated above (Fig. S1).

### DNA extraction from volcanic soils

0.25 g of soil was homogenized using a mortar and pestle, with cleaning using 70% (v/v) ethanol and distilled water between uses. DNA was extracted from homogenized soils using a DNeasy PowerSoil Pro Kit (Qiagen), with extracted DNA immediately cleaned using a PowerClean pro Cleanup Kit (Qiagen). DNA concentrations were calculated using a Qubit dsDNA HS Assay Kit (Thermo Fisher Scientific).

### Bacterial community composition

The 16S rRNA gene was amplified from ≥200 ng total DNA from each soil sample and sequenced using an Illumina PE250 sequencer at Novogene (Cambridge, UK). Sequences were quality-filtered using QIIME, including removal of low-quality reads and trimming based on Phred quality scores, before downstream processing (Caporaso *et al.,* 2010). For every sample, chimeras were removed using reference-based chimera checking with VSEARCH 2.16.0 (Rognes et al., 2016). The filtered reads were then clustered into operational taxonomic units (OTUs) at 97% similarity using UPARSE (Edgar 2013). Taxonomy was assigned using SILVA_v138 database (Quast *et al.,* 2013).

### Statistical analyses and OTU classification

Microbial alpha-diversity was expressed as species richness, Shannon index, species evenness and Simpson’s diversity. We used linear models to assess the effect of soil age (soils destroyed from the eruption at different times; 3-levels) on CO consumption rates and all four measures of alpha-diversity. For beta-diversity, we first normalised the OTU dataset using the Hellinger transformation (Legendre and Gallagher, 2001) with the ‘decostand’ function. Then we carried out principal coordinates analysis (PCoA) ordination on normalised data using the ‘pcoa’ function. All analyses were performed in R 4.3.2 using nlme, vegan, and ape packages.

Statistically significant differences between conditions for CO consumption tests were compared in triplicate by one-way analysis of variance (ANOVA) followed by a Tukey post hoc test using R studio using the vegan package.

### Metagenomic functional analysis

Total DNA (≥200 ng) for metagenomic analysis was also sequenced by Novogene (Cambridge, UK). FastQC (version 0.11.9) was employed to assess the quality of raw sequencing data obtained from the samples (Andrews 2019). Low-quality reads were trimmed using BBduk (version 38.86, quality-trim both ends to a phred score ≥ 15) (Bushnell, et al., 2017). Metagenomic assembly was performed to reconstruct the microbial genomes present in the environmental samples from the sequenced reads using SPAdes (version 3.14.0, Bankevich *et al.,* 2012; Nurk *et al.,* 2017) with Phred+33 quality score encoding (--phred-offset 33). Binning analysis was performed with scaffolds larger than 1000 bp. MetaWRAP (version 1.2.1) was used for binning analysis, leveraging three binning software programs: MaxBin2, MetaBAT2, and CONCOCT (Uritskiy *et al.,* 2018; Wu *et al.,* 2016; Kang *et al.,* 2015, Alneberg *et al.,* 2014). The resulting bins were further refined using the bin_refinement module within MetaWRAP, retaining only those with completeness >60% and contamination <10%. Identification was carried out using GTBDK database (GTDB release 10; R10-RS226) (Parks et al., 2025). Read mapping and genome coverage estimation of MAGs were performed using CoverM (v0.7.0) with the bwa-mem mapper on paired-end reads using default criteria (Aroney et al., 2025). To further enhance the characterization of the metagenomic data, BLASTx analysis (with an E-value threshold of 1×10⁻⁵) was used for comparing the *coxL* gene (form I characterized by the inferred amino acid motif AYXCSFR (Dunfield and King 2004) against our CoxL form I database. Further annotation was performed on MAGs containing *coxL* genes in their genome. For this, RAST (Rapid Annotation using Subsystem Technology) was employed for functional annotation using default parameters (Aziz, et al., 2008), focusing on CODH-related genes, including both form I and form II. A phylogenetic tree of the MAGs was inferred from the concatenated alignment of 120 bacterial marker genes using GTDB-Tk (version 2.5.2). The resulting unrooted tree was converted to an iTOL-compatible format for visualization and annotation, enabling the generation of a chiplot diagram to display phylogenetic relationships among the MAGs. The phylogenetic tree, along with an overall summary of the metagenomic analysis, including *coxL* gene identification, was built using Chiplot (Xie et al., 2023). Metagenomic analyses were performed on a high-performance computing cluster supported by the Research and Specialist Computing Support Service at the University of East Anglia (Norwich, UK).

## Results

### Soil physicochemical properties

Soil pH was very similar between the two oldest soils (PDB from the 1401 lava flow and ML from the 1559 lava flow) with averages of 5.2 and 5.3, respectively, while the most recent volcanic deposit had a near-neutral pH of 6.8 (Table S1). The youngest deposit (CDL from the 2007 lava flow) was substantially drier than the older samples with average soil moisture of 5.6 %, compared to 59.0 % and 64.7 % for the 1559 and 1401 samples, respectively, with the highest-moisture soil coinciding with the presence of vegetation (Table S1). Soil temperatures varied inconsistently among sites (20.2–28.2 °C), likely influenced more by local factors such as elevation and vegetation cover.

### CO uptake by soil microbes

All soil samples showed CO consumption occurring with no observable lag (Fig. 2) and this can be put down to biological activity due to no CO consumption observed in autoclaved samples (Fig. S1). CO consumption rates yielded significant variation among the sites (P<0.01). We observed that the oldest site (PDB, lava 1401) differed significantly from the other two sample sites (ML and CDL) with a faster rate of CO consumption of 0.63 nmol/h/g soil (Fig. 2). The 1559 (ML) and 2007 (CDL) sites showed much slower CO consumption rates and no significant difference between the two with 0.09 nmol/h/g soil and 0.06 nmol/h/g, respectively.

**Figure 2.**
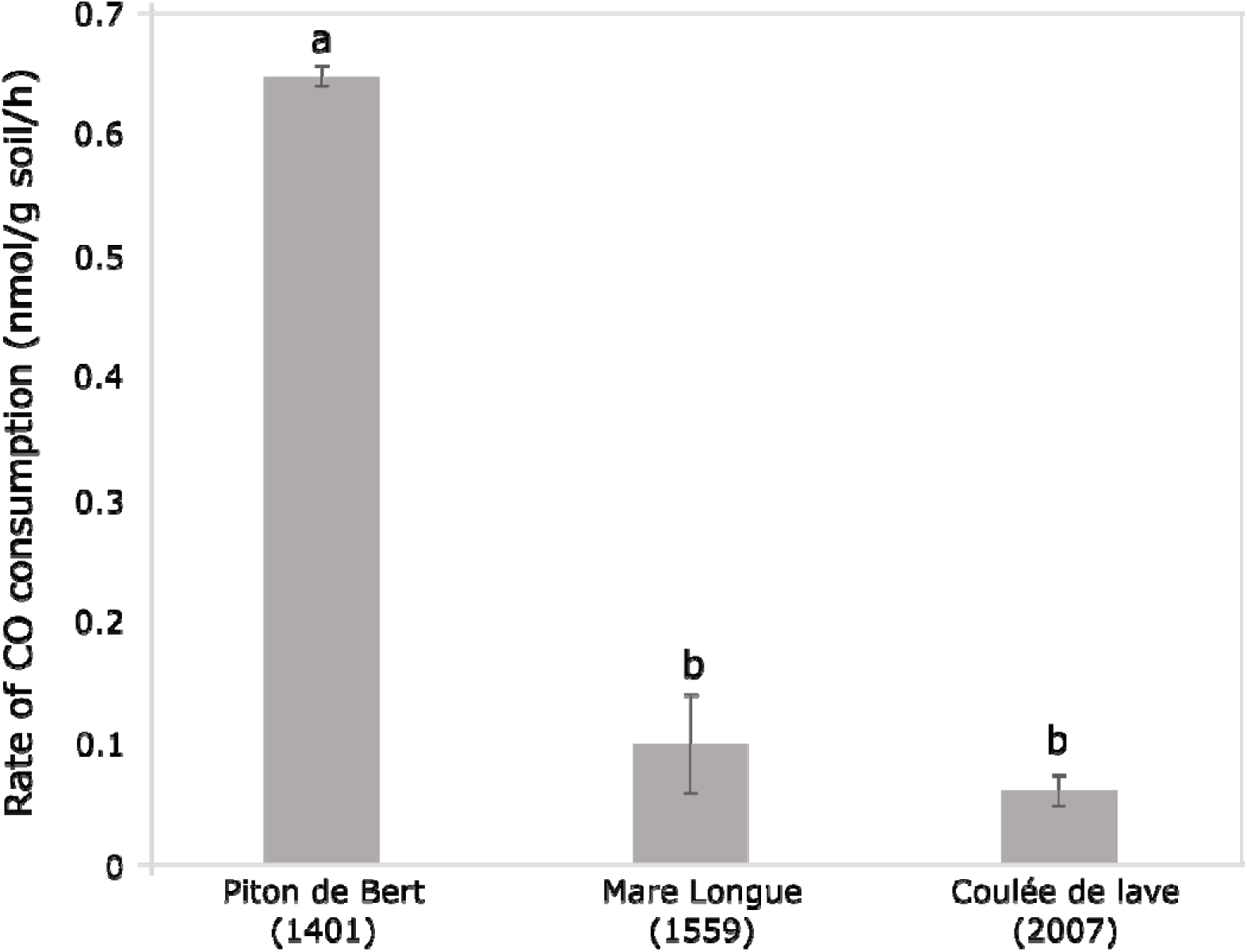
Consumption rates of CO (mean ± SD) in soils from three different locations on Piton de la Fournaise volcano in Réunion Island. Lower-case letters (a, b) denote differences in pairwise comparisons (P<0.05, Tukey’s HSD).

### Total microbial community composition

PCoA of bacterial OTUs showed differences in community composition for all three eruption sites due to separate clusters for all sampling sites (Fig. 3). Several notable patterns emerged from the relative abundance at the phylum level between the three sample sites (Fig. 4A). The phylum Acidobacteriota shows a steady increase in abundance from 10.78 % in the 2007 eruption sites (CDL) to 13.88 % in the 1559 eruption sites (ML) and then a substantial rise to 20.13 % in the 1401 eruption sites (PDB), indicating a progressive dominance over time. Actinomycetota, on the other hand, fluctuates inconsistently, with a decrease from 17.04 % in the 2007 eruption sites (CDL) to 10.42 % in the 1559 eruption sites (ML) and 9.90 % in the 1401 eruption sites (PDB). Bacteroidota, while maintaining a relatively low percentage overall, exhibits a decrease from 3.01 % in the 2007 eruption sites (CDL) to 0.3 5% in the 1401 eruption sites (PDB). In contrast, Pseudomonadota remains relatively stable, maintaining its dominance with percentages around 28-29 % across all three sample sites. Additionally, Chloroflexota demonstrates an increase from 2.35 % in the 2007 eruption site (CDL) to 4.25 % in the 1559 eruption site (ML), followed by a further rise to 4.34 % in the 1401 eruption site (PDB). Whereas Bacillota display a decrease from 1.42 % in the 2007 eruption sites (CDL) to 1.08 % in the 1401 eruption sites (PDB). Finally, Verrucomicrobiota, while initially present at 1.22 % in the 2007 eruption sites (CDL), experiences fluctuations, reaching 4.25 % in the 1559 eruption sites (ML) before declining to 2.61 % in the 1401 eruption sites (PDB).

**Figure 3.**
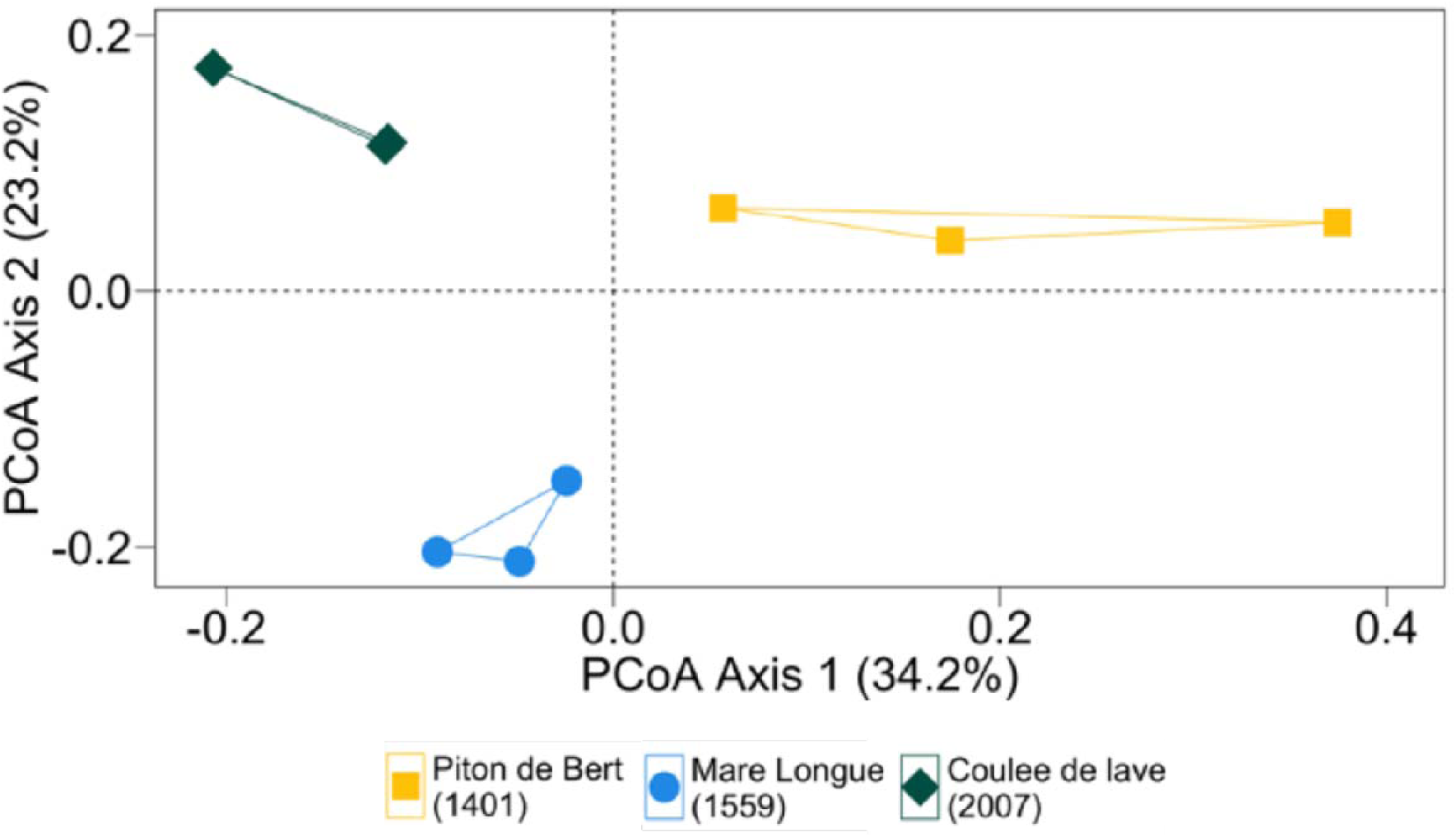
Principle coordinate analysis (PCoA) plots of OTUs (97% sequence similarity) derived from 16S rRNA genes extracted from soil. The legend indicates the eruption date of each sample.

**Figure 4.**
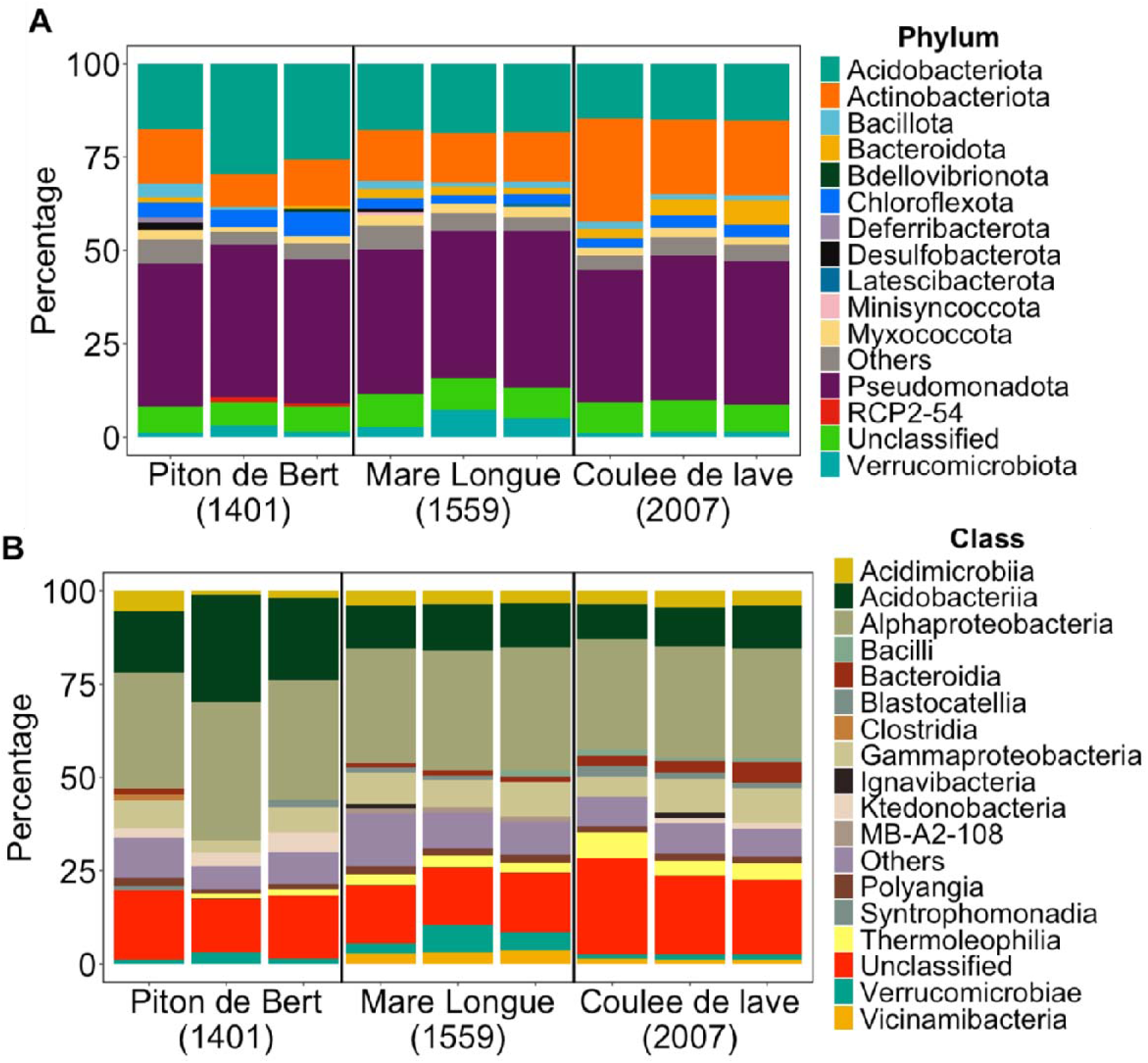
Relative abundance of microbial communities at the phylum (A) and class (B) level based on 16S rRNA genes in the different sample sites. “Unclassified” taxa are those OTUs that were not classified at the genus level. “Others” are those OTUs that were classified but the total abundance was less than 0.8% of all OTUs.

The composition of bacterial classes across the sample sites also displays variations (Fig. 4B). In the 2007 eruption sites (CDL), Alphaproteobacteria dominated at 29.62 %, followed by unidentified Actinobacteria at 13.19 % and Acidobacteriia at 10.80 %; however, in the 1559 eruption sites (ML), Alphaproteobacteria remained prevalent but slightly increased to 31.60 %, while Acidobacteriia decreased to 11.59 %. Contrastingly, in the 1401 eruption sites (PDB), Alphaproteobacteria maintained dominance but with a substantial increase to 33.05 %, accompanied by a significant rise in Acidobacteriia to 22.40 %. Additionally, there was a notable emergence of Ktenobacteria at 5.52%, which was not present in significant proportions in the 2007 (CDL) and 1559 (ML) eruption sites (Fig. 4B).

While all three sites exhibit high species diversity and evenness, there are discernible differences in these metrics between each site (Fig. 5). The 2007 eruption sites (CDL) stand out with the highest species richness and Shannon diversity index, indicating a richer and more diverse microbial community compared to the 1559 (ML) and 1401 (PDB) eruptions sites. However, all three sites demonstrate high Simpson indices, suggesting a similar level of species evenness across the sampled microbial communities.

**Figure 5.**
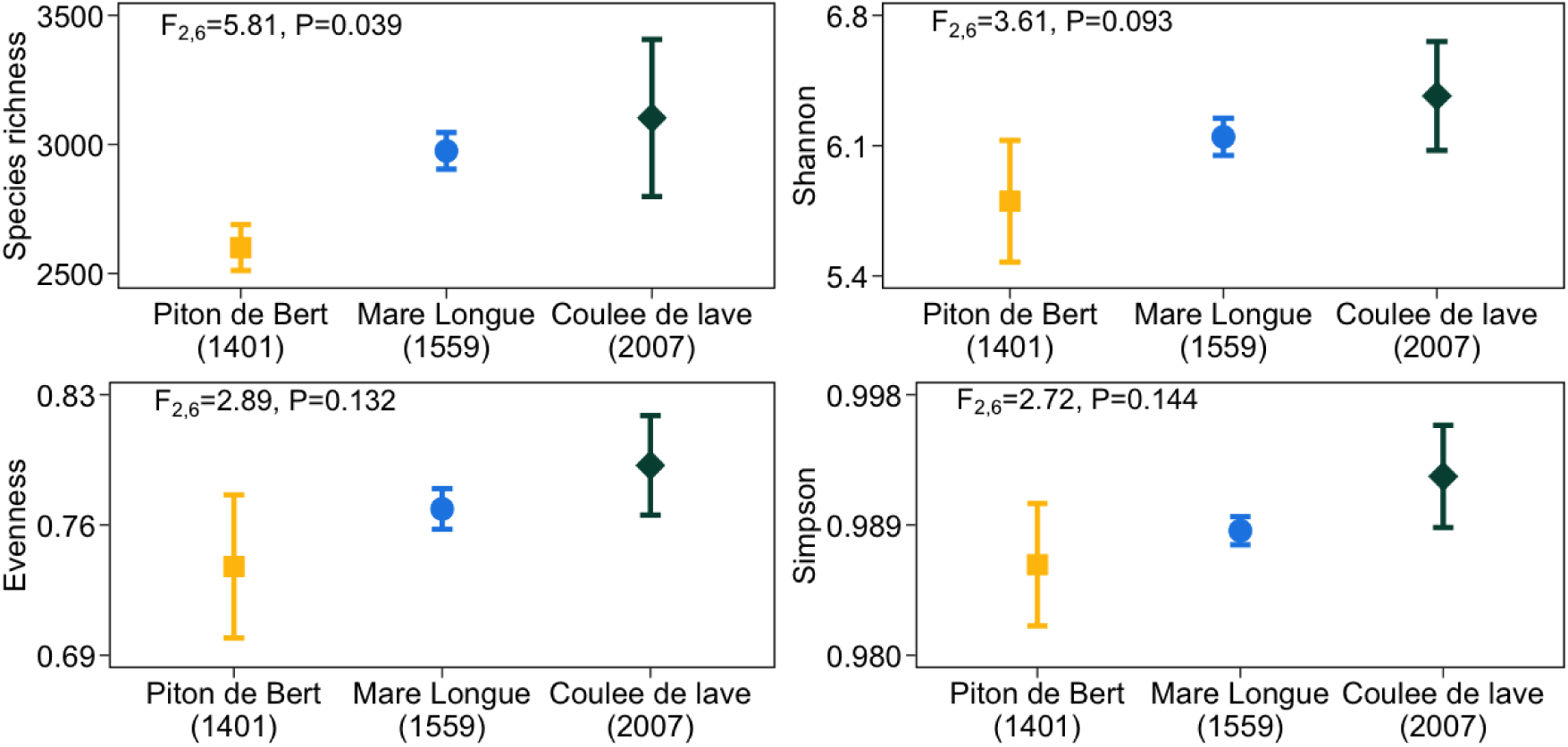
Alpha-diversity of microbes in soils from three different lava flows on Piton de la Fournaise volcano in Réunion Island. Alpha-diversity is evaluated as species richness, Shannon index, species evenness and Simpson’s diversity.

### Metagenomic analysis

A total of 13-14 million scaffold reads were recovered from the soil metagenomes for each site. Even though all the sites underwent similar sequencing efforts, the youngest soil (CDL from the 2007 lava flow) had the highest quality assembly, recovering a higher quantity of large scaffolds (308,753 scaffolds >1 kb) compared to the other sites (139,139 scaffolds >1 kb in the ML site, lava 1559 and 279,277 scaffolds >1 kb in the PDB site, lava 1401). 46 MAGs were retrieved in total. For our study, we analyzed only MAGs containing at least one copy of form I *coxL* gene, totaling 17 MAGs. 10 out of the 12 MAGs recovered from the 2007 lava samples (CDL) had >70% completeness and <5% contamination, compared to three MAGs from the 1559 soil (ML) samples and two MAGs from the 1401 samples (PDB), which had >60% completeness and <10% contamination. MAGs were affiliated to the phyla Acidobacteriota, Actinomycetota, Myxococcota, and Pseudomonadota (Fig. 6). In the oldest soil (PDB from the 1401 lava flow), environmental genomes were retrieved which related to the Pseudomonadota and the Actinomycetota. Environmental genomes from the 1559 soil (ML) were similarly related to the Actinomycetota and Pseudomonadota (Fig. 6). MAGs binned from the youngest soil (CDL from the 2007 lava flow) included three assigned to Pseudomonadota (family Xanthobacteraceae), two assigned to the Acidobacteriota, one Myxococcota, and six assigned to Actinomycetota (including the families Gaiellaceae, Ilumatobacteraceae, Solirubrobacteraceae, Streptosporangiaceae and Nocardioidaceae).

**Figure 6.**
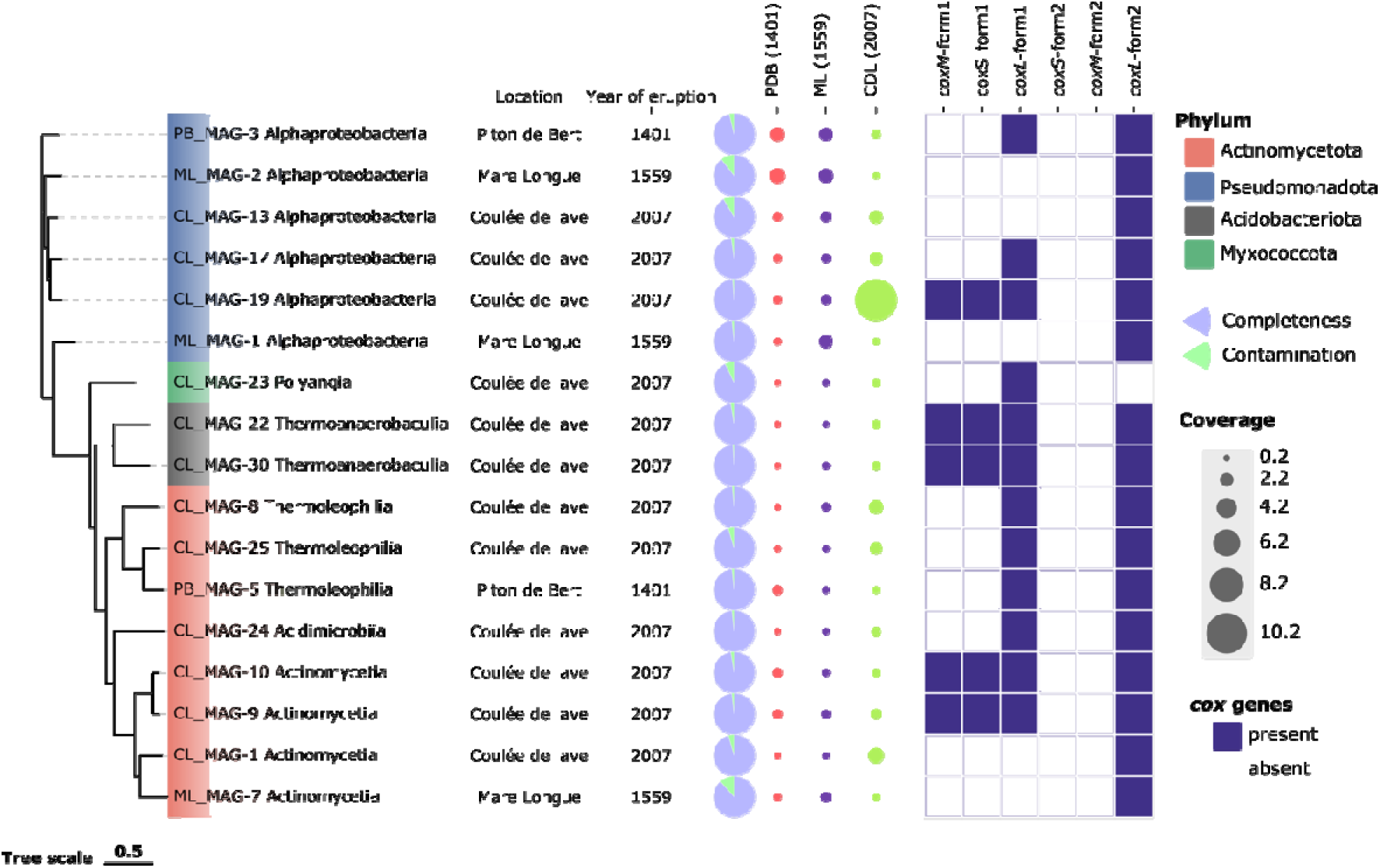
Analysis of MAGs retrieved from a chronosequence of volcanic deposits, classified according to completeness and contamination (pie charts), coverage, and the presence/absence of CODH-encoding genes.

Following BLASTx analysis, which confirmed that 17 out of the 46 MAGs initially retrieved had >50% form I *coxL* matches based on sequence identity, further annotation was performed on these MAGs. RAST was then employed for functional annotation, focusing on CODH-related genes, including both form I and form II. Our results showed that seven MAGs contained form I *coxL* alone, while a further five contained all three structural genes (*coxMSL*). 16 MAGs contained the putative form II *coxL* gene.

## Discussion

This study aimed to elucidate changes in microbial community composition, complexity, and CO oxidation rate in different aged volcanic deposits from Piton De La Fournaise Volcano. We identified clear differences in the relative abundance of known CO-oxidizing taxa in different lava flows, with the Pseudomonadota dominating in each site and the Ktedonobacteria, previously found to dominate in recent deposits (Hernandez *et al.,* 2020b), present at surprisingly low relative abundance in the younger site (CDL), which gradually increases with soil age (Fig. S1). Further, we found the oldest volcanic deposits to have the lowest microbial community diversity (Fig. 5) but the highest rates of CO oxidation (Fig. 2). Therefore, we have provided new evidence to suggest that CO oxidation remains an active metabolic process in older soils. However, we must reject the assumption that microbial community complexity increases with soil age, with other factors likely being greater drivers of increasing complexity in these environments.

The physicochemical properties observed in our study revealed notable trends with soil age (Table S1). Specifically, older soil samples exhibited lower pH levels, which could be attributed to the accumulation of organic matter over time. As organic matter decomposes, it generates organic acids that contribute to the acidity of the topsoil. Moreover, our findings indicate that soil moisture content increased with soil age. This may be linked to the enhanced soil structure resulting from the accumulation of organic matter, which allows for better water retention within the soil matrix (Chtouki et al., 2022).

The composition of the soil samples provided indirect evidence of the presence of vegetation, with older samples showing more visible signs of vegetation such as leaves and twigs compared to the predominantly rocky younger soil samples. This is as expected when looking at vegetation — specifically rainforest cover — on the island compared to our sample site locations (Strasberg 1994). This observation suggests a gradual transition to more habitable soil conditions over time, as older samples contained more vegetation material. This pattern could be interpreted as an early indication that biological complexity might increase with soil age, as has been reported in other volcanic systems (Gómez-Alvarez et al., 2006). However, as shown later in our results, this expected trend was not observed in our microbial data, suggesting that the relationship between vegetation development and microbial community change is not straightforward in this system.

The results obtained from the analysis of CO consumption across all different-aged volcanic soils provide compelling support for the hypothesis that CO-oxidizing microbes are active in younger soil samples. This is supported by consumption occurring in all samples with no observable lag (Fig. 2), as well as the absence of CO consumption in autoclaved samples indicating a biological process, distinguishing the role of living microbes from potential abiotic factors that could affect CO levels (Fig. S1). The previously discovered links between vegetation influence and the abundance and diversity of CO oxidizers might explain the observed differences in CO consumption rates, considering that older lava flows would have had more time for ecological succession, leading to increased organic matter and, potentially, a more developed microbial community (Weber and King 2010). An alternative explanation for the considerable increase in CO consumption in the older samples can be seen when comparing with the relative diversity at the class level. Another explanation could be that higher soil CO₂ concentrations promote may increase consumption. For instance, Bénard et al. (2023) found a higher rate of magmatic CO₂ release in soils at higher elevations on the island and that the magnitude of these effects and the type of transfer depend on the geological properties. The relative abundance of Ktedonobacteria rises significantly in the oldest 1401 (Fig. S2) site compared to the two younger sites, which matches the significant increase in carbon monoxide consumption, consistent with this previously being identified as a class with the genetic potential for CO oxidation (Hernández *et al.,* 2020a). These data demonstrate that despite minor differences in most community complexity indices between sites, the functional capacity of each community differed markedly in terms of CO oxidation, potentially reflecting changes in the composition and behavior of the resident microbial communities.

Using 16S rRNA gene sequencing data, our research offers an in-depth analysis of the microbial community structures found at various volcanic eruption sites on Réunion Island. We observed clear spatial segregation of microbial communities (Fig. 3), which corresponds directly to the lava flow’s age, suggesting a distinct community evolution over time. When using the data to classify organisms, all sites showed communities of bacteria present, confirming that these harsh environments can sustain microbial life. At the phylum and class levels, our data indicate significant variations between sites (Fig. 4). For example, older sites like the 1401 Piton de Bert showed a greater abundance of Acidobacteriota than in the newer sites, with a similar rise in Acidobacteriia classes at the oldest site further supporting this trend (Fig. 4B). Conversely, Actinomycetota does not exhibit a consistent pattern, decreasing with the site’s age, which implies that other environmental factors may impact certain microbial taxa leading to them being outcompeted. This underscores the dynamic nature of microbial community assembly and ecological succession. Interestingly, Pseudomonadota consistently dominates across all sites, likely due to their capacity for utilizing diverse substrates and rapidly colonizing new areas, playing a crucial role in primary ecological succession (Zhou *et al.,* 2020). Studies have previously found the Pseudomonadota, a group that contains diverse characterized CO-oxidizing bacteria (Santiago *et al.,* 1999; Dawson *et al.,* 2025), are dominant in certain vegetated volcanic deposits (Weber and King 2010; Hernández *et al.,* 2020b) or unvegetated deposits across diverse environments (Byloos *et al*., 2018).

When analyzing community species richness and evenness, the highest species richness and Shannon diversity indices occur at the youngest site (Fig. 5), contrasting with the hypothesis that microbial community complexity increases with soil age. This suggests that these sites experience rapid colonization by diverse species immediately following an eruption, with drivers of diversity such as associated vegetation potentially playing a greater role than the age of each deposit. Over time, as ecological succession progresses, the community composition becomes more evenly structured, as evidenced by high Simpson indices at all sites. This is evident at the class level where there is a notable decline of various classes such as Ignavibacteria and Bacilli in the older sites (Fig. 4B), which is likely due to competition from other classes that benefit more from the shift in available nutrients provided by nutrient cycling due to this diverse initial microbial colonization. For example, Ignavibacteria, facultative or strictly anaerobes which have previously been found in microbial mats (Iino *et al.,* 2010), might be more prevalent in newer volcanic soils before aerobic microbes like Alphaproteobacteria establish dominance. Additionally, the presence of Bdellovibrionota in the oldest sample site is interesting due to it having been found previously to be an obligate predator which can consume other Gram-negative bacteria, so this group could potentially be playing a role in the declining diversity (Li *et al.,* 2021). Comparing our findings with prior research, such as studies conducted on the Kilauea Volcano in Hawaii, shows similarities in dominant phyla across different volcanic sites (Gomez-Alvarez *et al.,* 2007). However, other global volcanic studies, including Llaima Volcano in Chile, often find younger soils with lower diversity indices and fewer species, which tend to increase as the soil matures and stabilizes due to complex ecological interactions (King *et al.,* 2003b; Gomez-Alvarez *et al.,* 2007; Hernández *et al.,* 2020b). Our results, particularly the high species richness in the youngest soils, differ from these findings, suggesting that initial colonization dynamics may also be influenced by site-specific factors such as elevation in addition to soil age.

Our study provides evidence supporting the hypothesis that microbes capable of oxidizing carbon monoxide are present in volcanic soils, but CO consumption activity was highest at the oldest site. For example, bacterial phyla such as the alphaproteobacteria, which dominates all soil sample sites, have been previously identified to contain the CODH-encoding gene cluster (King 2003b; Hernández *et al.,* 2020a; Dunfield *et al.,* 2004; Schübel *et al.,* 1995). Additionally, MAGs which contained all genes required to express CODH (*coxMSL*) were exclusively isolated from the youngest soil (CDL) (Fig. 6). Ktedonobacteria, which has had extensive research into its CODH-encoding gene cluster through metagenomic studies and isolate-based studies (Hernández *et al.,* 2020a; King and King 2014; Islam *et al.,* 2019), showed an increase in relative abundance in the oldest 1401 sample compared to the two younger soil sites (Figure S2). This indicates that these CO-oxidising bacteria are not only essential in the initial colonization of these extreme environments, but also remain prevalent even when organic carbon content is not as limited, potentially showing that CO energy metabolism could be advantageous in these sites for quite some time. A study of microbial respiration in soils on Kilauea Volcano (King 2003a) also suggested that CO oxidation was selected for by microbial communities even when heterotrophic metabolism was possible.

In the exploration of soil metagenomes across different eruption ages, the recovery of MAGs demonstrated a notable disparity influenced by soil age. The most comprehensive assembly was achieved in the youngest soil (CDL), which boasted the highest number of large scaffolds, suggesting robust assembly quality. This superior assembly quality and more effective binning of genomic data in the youngest soil could be attributed to less genomic complexity at this stage of soil development, which would be in line with the hypothesis of complexity increasing with age from eruption; however, that challenges the previous results from analysis of 16S rRNA sequencing data. The taxonomic profiling of the MAGs revealed distinct phyla predominance corresponding to soil age. In the youngest soil, one quarter of MAGs were affiliated with the Pseudomonadota, including families known for their versatile metabolic capabilities, which may confer advantages in early soil colonization stages (Dawson *et al.,* 2025). Two MAGs recovered from the middle-aged soil were also affiliated with the Pseudomonadota, while the third was affiliated with the Actinomycetota, which is known to rely more on degrading complex organic compounds, potentially reflecting a shift in the organic composition of the soil as it matures. The oldest soil, similarly, yielded two MAGs related to Pseudomonadota and the Actinomycetota (Fig. 6), which may similarly reflect a shift to more metabolically versatile microorganisms.

### Conclusions

In conclusion, while our study did not fully support the initial hypothesis regarding an increase in microbial community complexity with increasing soil age, it did enhance our understanding of the role of CO oxidizing microbes in volcanic soils. Future research should focus on the isolation of key microbial strains to further understand their roles and contributions to ecological succession and soil development in extreme environments. Overall, our research contributes to the growing evidence that volcanic eruption sites serve as unique natural laboratories for examining microbial colonization and succession. The initial high diversity phase immediately following an eruption, coupled with subsequent microbial community stabilization, highlights the complex and dynamic nature of these ecosystems. These findings emphasize the need for further longitudinal and biogeochemical studies to fully understand the environmental factors and temporal dynamics influencing microbial communities in volcanic landscapes, which will enhance our predictions of ecological recovery and inform conservation strategies in extreme environments.

## Acknowledgements

We acknowledge the Doctoral School of Earth Sciences and Ecology of the University of Tartu for organizing the research expedition to Réunion island. The authors would like to thank the Observatory of the Science of the Universe-Réunion (OSU-R) for access to the facilities at the Mare Longue Research Field Station during the field campaign and acknowledge the National Park of La Réunion for issuing the collecting permit DIR-I-2022-281.

## Data Availability

Sequence data were deposited in the NCBI Sequence Read Archive (SRA) under the bioproject accession numbers: PRJNA1367882 for amplicon-sequencing data and PRJNA1367899 for metagenome-assembled genomes.

## Author contributions

MH conceived the project and secured the funding; ME and CAP contributed to the collection of the samples; CW and RAD performed the experiments; CW and SR performed 16S rRNA gene amplicon sequence analysis; MH performed metagenomic analysis. CW and MH drafted the paper; all authors contributed to its revisions and approved the final version.

