## Supplementary Information for "Distribution of carbon monoxide-oxidizing microorganisms along a chronosequence on Piton De La Fournaise volcano"

Table S1
Figure S1
Figure S2

Table S1. Location and physico-chemical properties of the soil samples

| Site | Coordinates | Eruption date | Average pH | Average soil moisture (%) | Soil temp ($^oC$) | Air temp ($^oC$) | Soil observations |
|---|---|---|---|---|---|---|---|
| Piton de Bert (PDB) | 21.2788831 S, 55.6980607 E | 1401 | 5.2±0.3 | 64.7 | 28.2 | 33.7 | Damp, no rocks, some vegetation |
| Mare Longue (ML) | 21.3512651 S, 55.7392276 E | 1559 | 5.3±0.1 | 59.0 | 20.2 | 21.3 | Dry, lots of rocks |
| Coulée de Lave (CDL) | 21.2866284 S, 55.7957900 E | 2007 | 6.8±0.3 | 5.6 | 25.1 | 29.5 | Dry, lots of rocks |

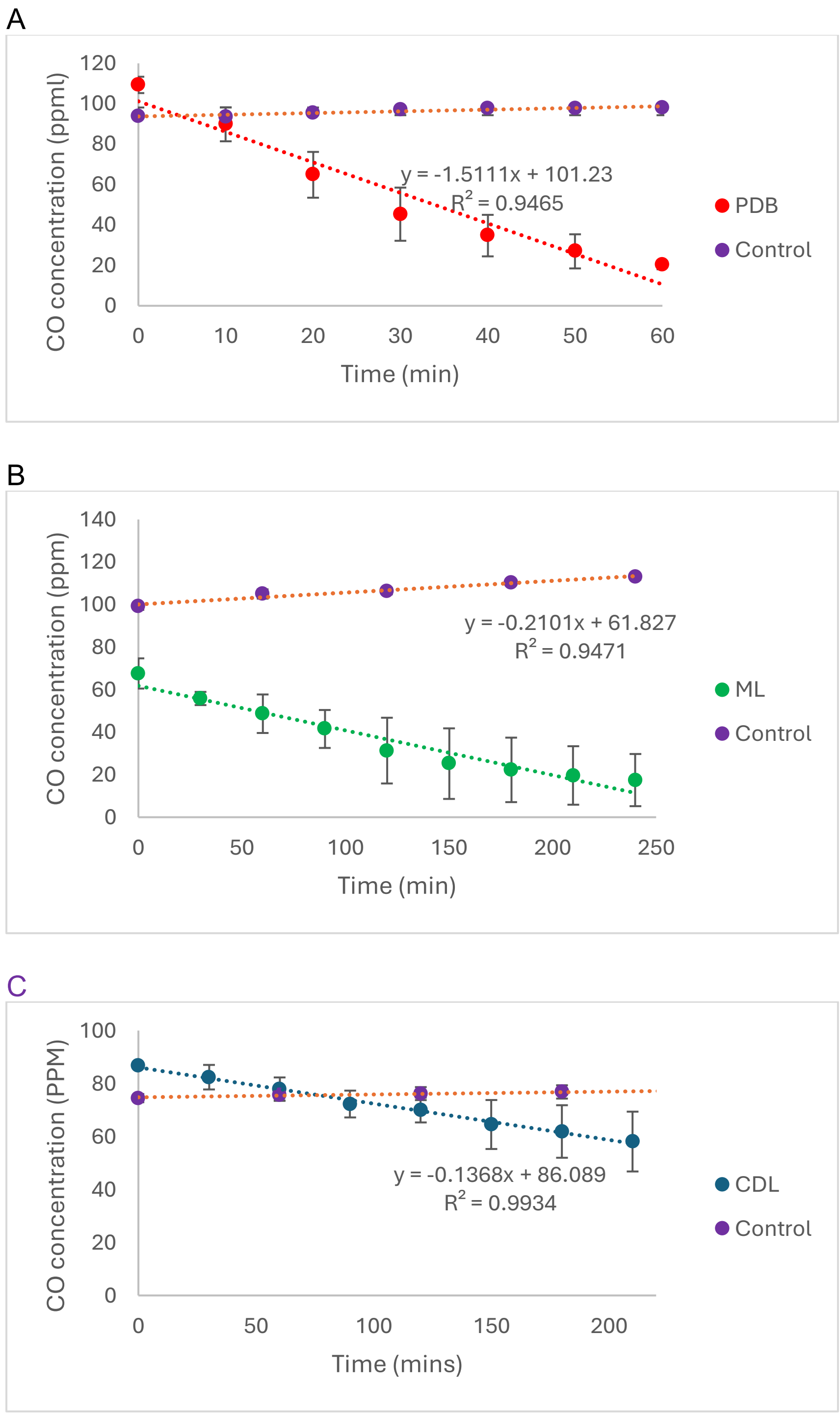


Figure S1. Rate of CO consumption by Piton de Bert (PDB) (A), Mare Longue (ML) (B) and Coulée de lave (CDL) (C) soil samples compared to controls from autoclaves soil samples. Points represent mean values with standard deviations of independent triplicate incubations.

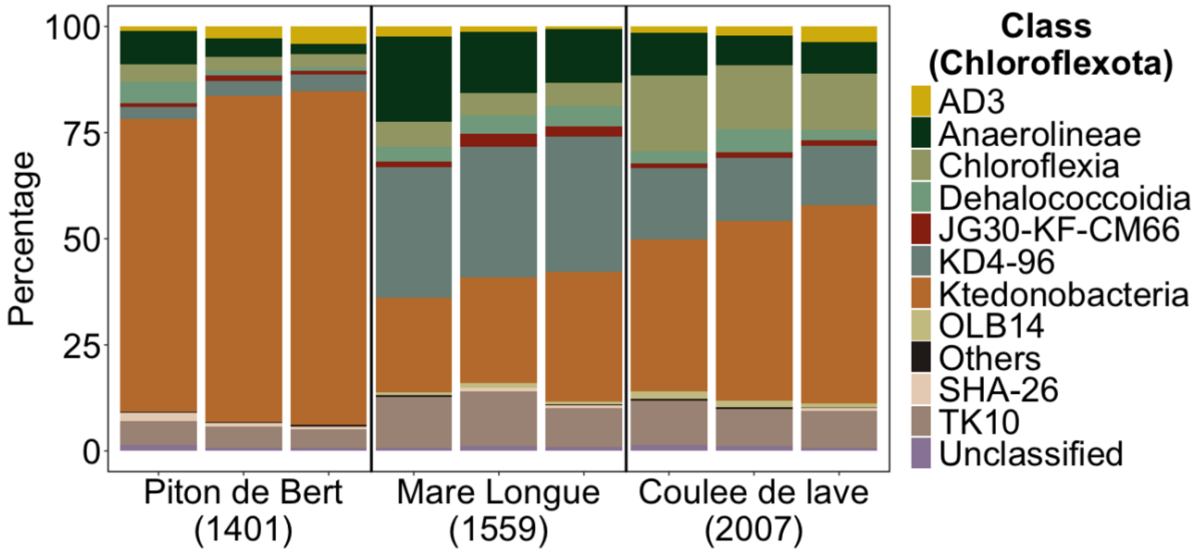


Figure S2. Relative abundance of microbial communities at the Chloroflexota class level based on 16S rRNA genes in the different sample sites. “Unclassified” taxa are those OTUs that were not classified at the genus level. “Others” are those OTUs that were classified but the total abundance was less than 0.5% of all OTUs.
